# Transcription factor-target relationships complicated by knockout analysis

**DOI:** 10.1101/2020.08.30.274548

**Authors:** Zhiming Dai

## Abstract

Knockout analysis is a common tool to reveal transcription factor (TF) functions. However, such a reverse genetic analysis based on observed phenotype changes in mutant cell may lead to a misunderstanding of TF wild-type functions. Here, a model was proposed, in which the knockout-observed TF-target regulatory relationships might only occur in mutant cell, and they do not reflect TF normal functions in wild-type cell. Actually, the knockout of one TF might release another TF which is the protein-protein interaction partner of the deleted TF. The free TF could bind its new target genes and cause their significant expression changes. These seemingly TF knockout affected genes are thus not directly regulated by the deleted TF, but are gain-of-regulated genes of the latter TF in mutant cell. Based on this model, multiple sources of genome-wide data were used to identify 20 such TF pairs, and one pair was validated using other independent data. TF wild-type regulatory genes are not associated with their gain-of-regulated genes. My findings revealed TF-target relationships complicated by TF knockout analysis.

## Introduction

Genetic analysis is one basic research strategy to understand genes of interest [1]. A common application is to measure expression changes in response to perturbation of one gene [2-4]. With the assumption that the perturbed gene is likely to be responsible for the changes, the cellular function of this gene in the wild-type cell may be revealed by a reverse genetic analysis [5, 6].

However, perturbation experiments may have some limitations. First, it generally infers wild-type functions of one gene from the phenotype changes caused by perturbation of this gene. The revealed cellular functions may be only perturbation-specific, that is, this reverse genetic analysis may not reflect the normal functions of this gene in the wild-type cell. The gap between wild-type cell and perturbation cell may lead to a misunderstanding of the living system [7]. Second, it does not generally provide information about whether the perturbation of one gene causes phenotype changes directly or indirectly. Moreover, perturbation experiments alone may be unable to reveal the actual genes that directly cause phenotype changes. Knockout analysis is one powerful genetic analysis tool to reveal cellular function of one gene of interest. Over the past few years, such experiments on transcription factor (TF) knockout helped identify TF potential regulatory targets [8-11]. TF perturbation experiments serve as complementary data sources to ChIP-chip (or ChIP-seq) experiments to uncover TF cellular functions [12-14]. Several large-scale TF perturbation datasets are now available [8, 9]. A compendium of 269 TF knockouts in budding yeast was published by Hu et al., and the differentially expressed gene targets were compared with the TF direct bound gene targets measured by ChIP-chip [15]. Only ∼3% of TF knockout affected genes are bound by the corresponding TF [16], indicating that TF indirect regulation through regulatory cascades may be prevalent. However, removal of indirect TFs-target regulatory interactions resulted in negligible improvements to the overlap [15]. The TF knockout expression data has been reanalyzed by Reimand et al. and more TF knockout affected genes were identified that are bound by the corresponding TF, but the overlap is still very small [17]. Note that these TF-DNA binding data were measured in wild-type cells, whereas TF knockout affected genes were identified in TF-deficient cells.

In this study, one possible explanation was hypothesized for the small overlap between TF knockout affected genes and TF bound genes is that TF knockout analysis may not reveal some TF native functions in wild-type cell. If one specific TF A does not regulate one specific gene (through direct TF-DNA binding or regulatory cascades) in wild-type cell, why the expression of this gene shows significant changes upon the knockout of TF A? One possibility is that the knockout of TF A leads to a rewiring of transcriptional regulatory program of this gene. Knockout of TF A was assumed to release another TF B, which is protein-protein interaction (PPI) partner of TF A in wild-type cell. The free TF B may bind its DNA motif in this gene, leading to the apparent observation that this gene shows significant changes in gene expression in TF A-deficient mutant. Based on these assumptions, a model was proposed, in which knockout of one TF might cause expression change of one gene, but this change is not directly linked to this TF but linked to another TF gain-of-binding to this gene. Moreover, these two TFs have no regulatory roles on this gene in the wild-type cell. Here, 20 such TF pairs were identified in budding yeast using various sources of datasets and strict criteria. Further analysis of these TF pairs revealed that TF knockout might create TF functions that are irrelevant to their native functions.

## Materials and methods

### Data preparation

Genome-wide binding affinity data corresponding to 203 TFs in wild-type were taken from Harbison et al. [16]. A P value cutoff of 0.005 was used to define the set of genes bound by a particular TF. Combining several motif finding methods, DNA motifs that are not bound by the corresponding TFs in each promoter (an average size of 480 bp) in wild-type were also identified by Harbison et al. [16]. The presence of DNA motifs implies that the promoter may be bound by the corresponding TFs in other cellular conditions. Genome-wide changes in gene expression data corresponding to the knockout of 269 TFs were taken from Hu et al. [15]. Differentially expressed genes for each TF knockout were identified by Hu et al. [15] and Reimand et al. [17], respectively. As the identification method of Reimand et al. were reported to have improvement compared with that of Hu et al., the list of differentially expressed genes from Reimand et al. was used in this study. The TF knockout data and TF binding data have 188 TFs in common. For TF knockout affected genes of each TF, those genes that are bound genes of the corresponding TF were excluded. PPI data were taken from Stark et al. [18].

Genome-wide gene expression data used for co-expression analysis were measured under normal growth conditions [19-21], a total of 112 time points. For each potential TF, pair-wise Pearson expression correlation coefficient between its gain-of-regulated genes and its wild-type bound genes was calculated, pair-wise Pearson expression correlation coefficient between its gain-of-regulated genes and its knockout affected genes was calculated. For each apparent-potential TF pair, pair-wise Pearson expression correlation coefficient between potential TF gain-of-regulated genes and apparent TF wild-type bound genes was calculated. Pair-wise Pearson expression correlation coefficient between every possible gene pair was calculated.

### Method for identification of TF pairs

Whether there exist TF pairs in budding yeast as described in Figure 1 was examined. Integrating TF knockout data[15], TF binding data[16], TF binding motifs data[16] and PPI data[18], TF pairs meeting following criteria in the wild-type cell were identified (Figure 2A): (1) TF A shows PPI with TF B. (2) All TF A knockout affected genes are not statistically significantly bound by TF A or TF B (P>0.05, chi-square test). (3) All TF A knockout affected genes show statistically significant enrichment with motifs (not bound in wild type) of TF B (P < 10-5, chi-square test) but TF A (P>0.05, chi-square test). (4) TF A or TF B do not show PPI with TFs that have binding target enrichment (P<0.05, chi-square test) in all TF A knockout affected genes (i.e. TF A knockout affected genes are statistically significantly bound by the TFs). (5) All TF A knockout affected genes are not statistically significantly regulated by TF B (i.e. after knockout of TF B, these genes do not show statistically significant expression changes compared with the remaining genes (P>0.05, Mann-Whitney U-test)). In addition, the gene encoding TF B should not be regulatory target of TF A, that is, the gene encoding TF B is not differentially expressed gene for TF A knockout according to the list of differentially expressed genes from Reimand et al.. These strict criteria were used to control false positive. Together, it is less likely that either TF A or TF B regulate TF A knockout affected genes in the wild-type cell. Without the PPI partner TF A in the TF A-deficient cell, it is likely that the free TF B could bind TF A knockout affected genes which have DNA motifs for potential TF B binding, to cause their changes in gene expression.

**Figure 1.**
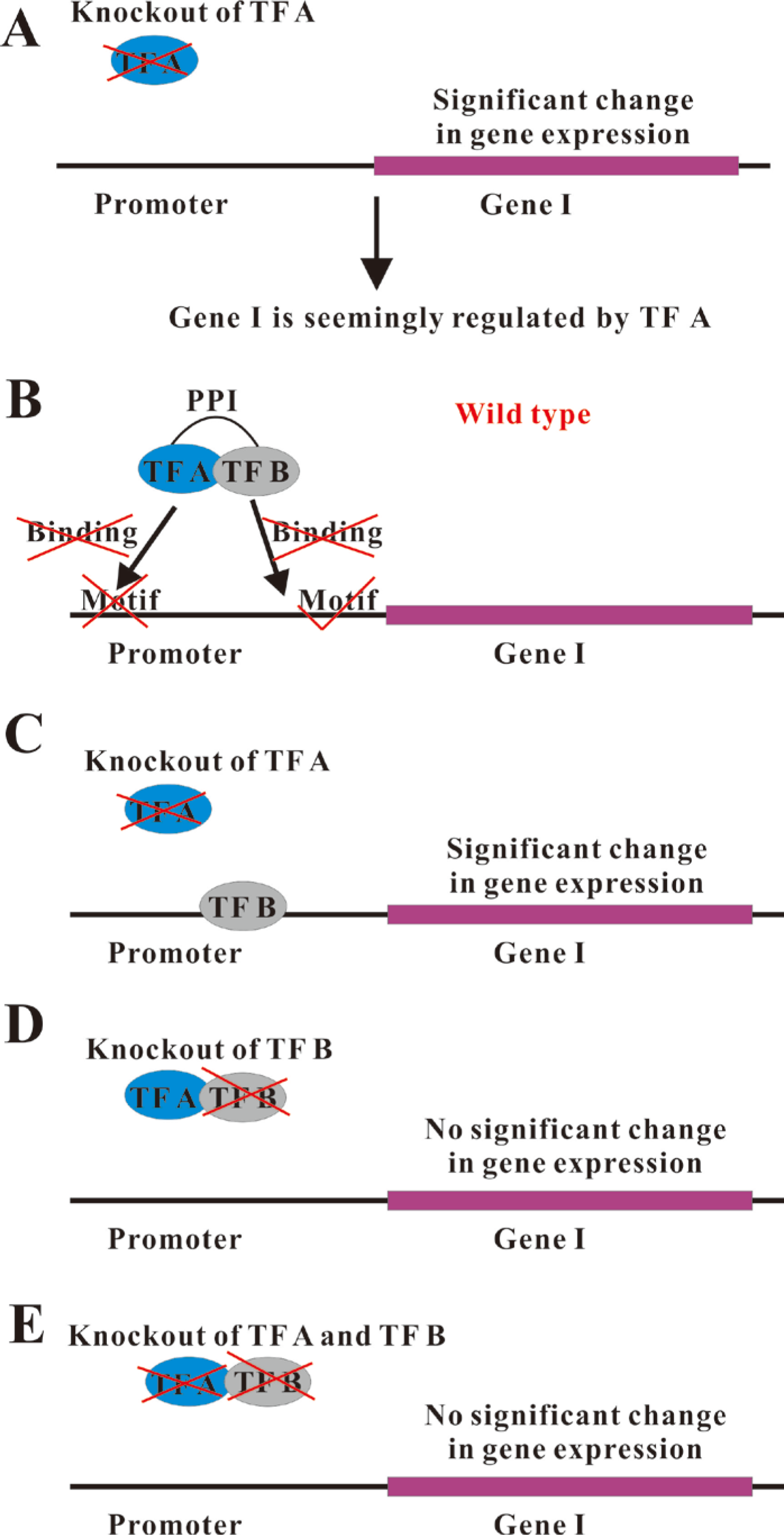
Transcription factor-target relationships complicated by knockout analysis. (A) Knockout of TF A can cause significant expression changes of Gene I. This gene is commonly described as a regulatory gene of TF A. (B) However, this so-called regulatory gene is not bound by TF A, and even have no TF A bound motifs in promoter region. Thus this gene may not be regulated by TF A in the wild-type cell. On the other hand, this gene has DNA motifs for TF B potential binding, though this gene is not bound by TF B in the wild-type cell. TF A and TF B have PPI. (C) Knockout of TF A may release its PPI partner TF B. The free TF B could find its bound motifs in Gene I, though TF B could not bind Gene I in wild type due to the limitation imposed by its PPI partner TF A. The so-called TF A regulatory gene is actually TF B gain-of-regulatory gene in TF A-deficient mutant. This actual regulatory relationship could not be revealed by knockout of TF A. Moreover, This gain-of-regulatory relationship could not be observed by either knockout of the TF B (D) or double knockouts of TF A and TF B (E).

**Figure 2.**
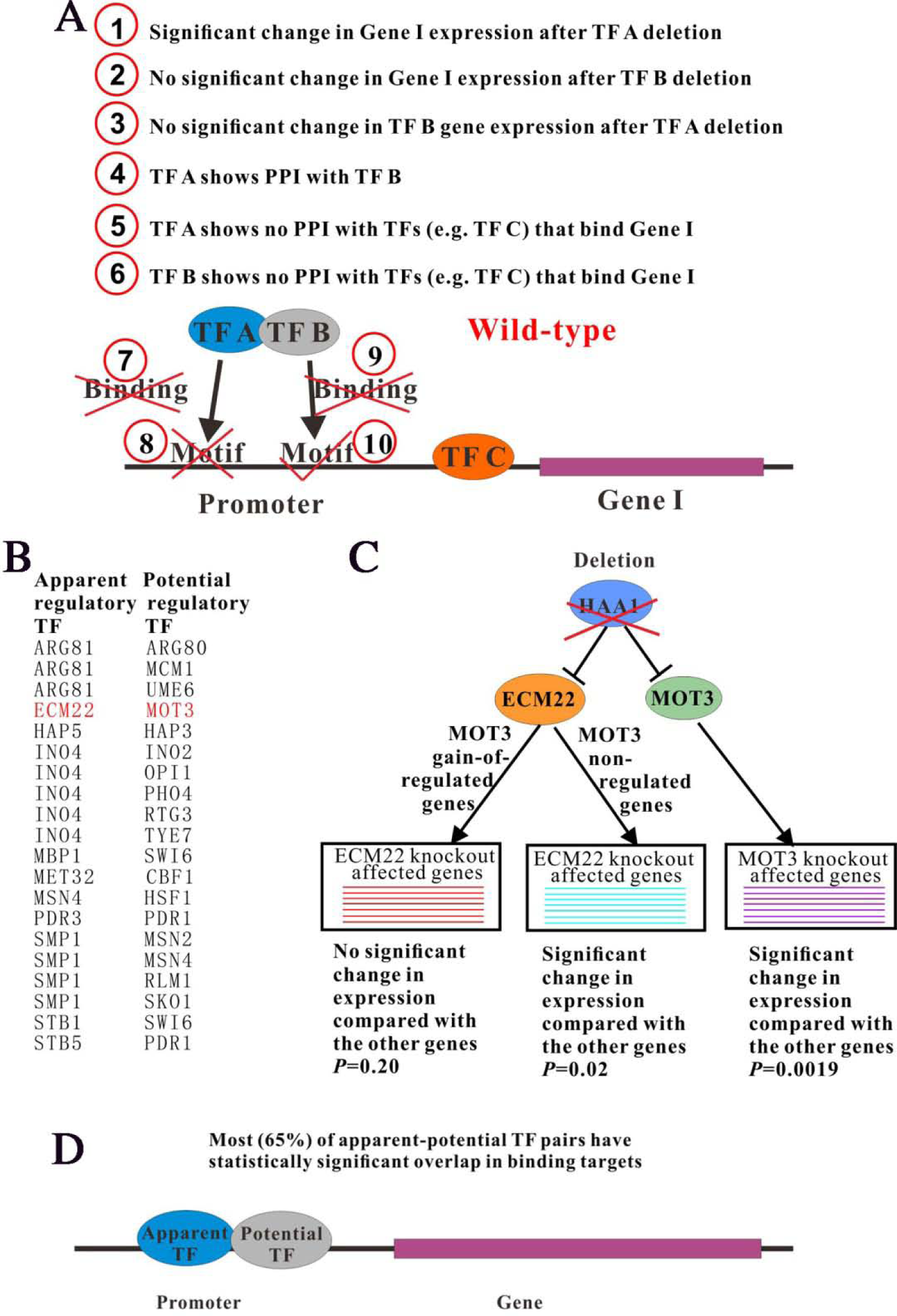
Identification of apparent-potential TF pairs described in Figure 1. (A) The criteria used to identify apparent-potential TF pairs. See method details in main text. (B) The list of identified apparent-potential TF pairs. The pair in red was validated in (C). (C) One validated TF pair. Genes encoding ECM22 and MOT3 were differentially down-expressed when the knockout of HAA1. Knockout of HAA1 was used as an approximation to double down-regulation of both ECM22 and MOT3. MOT3 knockout affected genes showed statistically significant expression changes compared with the remaining genes when HAA1 was deleted. MOT3 non-regulated genes in the ECM22-deficient cell showed statistically significant changes in gene expression compared with the remaining genes upon knockout of HAA1. If MOT3 regulated a portion of ECM22 knockout affected genes in the ECM22-deficient cell, knockout of HAA1 should reduce MOT3 regulatory effects. Indeed, MOT3 gain-of-regulated genes in the ECM22-deficient cell showed comparable changes in gene expression with the remaining genes upon knockout of HAA1. The statistical significant values calculated from Mann-Whitney U-test were indicated. (D) Most of apparent-potential TF pairs are cooperative TFs in wild type. 13 (65%) of 20 apparent-potential TF pairs bind target genes in common.

## Statistical Method

Chi-square test was used to examine whether a group of genes were statistically significantly bound by one TF. The number of genes in this group bound by the TF was considered as the observed number. The number of genes in this group was considered as the observed number. The number of all genes in budding yeast was considered as the expected number. The number of all genes bound by the TF was considered as the expected number. Chi-square test was used to examine the difference between the expected numbers and the observed numbers. Similarly, Chi-square test was also used to examine whether a group of genes show statistically significant enrichment with motifs (not bound in wild type) of one TF. The number of genes in this group having the motifs of the TF was considered as the observed number. The number of genes in this group was considered as the observed number. The number of all genes was considered as the expected number. The number of all genes having the motifs of the TF was considered as the expected number.

Given two samples of values, the Mann-Whitney U-test is designed to examine whether they have equal medians. The main advantage of this test is that it makes no assumption that the samples are from normal distributions.

## Results

### The proposed model

A model was proposed to reveal TF-target regulatory relationship complicated by knockout analysis. In this model, knockout of TF A can cause significant expression changes of Gene I (Figure 1A). This gene is commonly described as regulatory gene of TF A. However, this so-called regulatory gene is not bound by TF A, and even have no DNA motifs for TF A potential binding. Gene I thus may not be regulated by TF A in the wild-type cell (Figure 1B). This regulatory relationship may be specific only for TF A-deficient mutant. Moreover, this seemingly regulatory relationship may be only mediated by the knockout of TF A, it may not reveal the actual TF that is directly responsible for the expression changes of Gene I in TF A-deficient mutant.

Knockout of TF A may release its PPI partner (i.e. TF B). The free TF B could find its bound motifs in Gene I in TF A-deficient mutant, though the TF B could not bind Gene I in wild type due to the limitation imposed by its PPI partner TF A. Then this binding may cause differential expression of Gene I (Figure 1C). The so-called TF A regulatory gene (i.e. Gene I) is actually TF B gain-of-regulatory gene in TF A-deficient mutant. This actual regulatory relationship could not be revealed by knockout of TF A. On the other hand, as TF B does not bind Gene I in the wild-type cell, knockout of TF B causes no effect on TF binding and gene expression of Gene I (Figure 1D). Moreover, double knockouts of TF A and TF B may not alter TF binding and gene expression of Gene I (Figure 1E).

Together, though it is obvious that both TF A and TF B have no native regulatory functions on Gene I in the wild-type cell, Gene I is still identified as TF A knockout affected gene. In fact, Gene I is directly bound by TF B in the TF A-deficient cell. However, this relationship cannot be observed in the TF B-deficient cell and even in the TF A-TF B double-deficient cell. Thus, Gene I is generally not identified as TF B regulated gene. It is necessary to integrate other complementary data to reveal this phenomenon.

### Identification of TF pairs

Whether there exist such TF pairs described in the proposed model was examined by using strict criteria (Figure 2A, see details in Materials and Methods section). In this way, 20 TF pairs were identified (Figure 2B). These TF pairs were referred to as apparent-potential TF pairs, in which TF A and TF B are the apparent and potential TFs that might be responsible for gene expression changes of TF A knockout affected genes in the TF A-deficient cell, respectively. For each apparent-potential TF pair, apparent TF knockout affected genes meeting following criteria were identified: (1) They do not show significant expression changes (<2 fold changes) after knockout of potential TF. (2) They are not bound by apparent TF or potential TF in the wild-type cell. (3) They have DNA motifs that can be potentially bound by potential TF but apparent TF. These genes were referred to as potential TF gain-of-regulated genes in the apparent TF-deficient cell. On the other hand, apparent TF knockout affected genes meeting following criteria were identified: (1) They are not bound by potential TF in the wild-type cell. (2) They have no DNA motifs that can be bound by potential TF. These genes were referred to as potential TF non-regulated genes in the apparent TF-deficient cell. Note that both of gene groups are apparent TF knockout affected genes.

### One validated TF pair

Whether potential TFs indeed become gain-of-regulators of their paired apparent TF knockout affected genes after knockout of the apparent TFs was examined. If this is the case, double knockouts of one apparent-potential TF pair should not cause significant expression changes in apparent TF knockout affected genes that contain potential TF binding motif (i.e. gain-of-regulated genes of potential TF). Due to lack of double TF knockout data, single TF knockout data was used as an approximation. If one apparent-potential TF pair both show statistically significant down-expression when the knockout of another TF, this mutant could be used as an approximation to double knockouts of this apparent-potential TF pair. Using the list of differentially expressed genes from Reimand et al. [17], there are 3 such pairs among the 20 apparent-potential TF pairs: genes encoding ECM22 and MOT3 are differentially down-expressed when the knockout of HAA1 (i.e. up-regulated by HAA1), genes encoding SMP1 and MSN2 up-regulated by HMS1 and TYE7, genes encoding SMP1 and RLM1 up-regulated by HMS1 and TYE7. Next, whether this approximation is appropriate was examined. To address this issue, whether potential TF knockout affected genes are differentially expressed when the knockout of up-regulator of the potential TF was tested. Among the 3 apparent-potential TF pairs identified above, only one pair (ECM22-MOT3) demonstrated this property: MOT3 knockout affected genes show statistically significant expression changes compared with the other genes when the up-regulator HAA1 is deleted (P=0.0019, Mann-Whitney U-test, Figure 2C, S1). For the other 2 apparent-potential TF pairs (SMP1-MSN2 and SMP1-RLM1), the statistical significances were very weak (P>0.3, Mann-Whitney U-test). Thus the ECM22-MOT3 TF pair was focused on. ECM22 knockout affected genes were divided into two groups: MOT3 gain-of-regulated genes and MOT3 non-regulated genes in the ECM22-deficient cell. MOT3 non-regulated genes showed statistically significant changes in gene expression compared with the other genes upon knockout of HAA1 (P=0.02, Mann-Whitney U-test, Figure 2C). To sum up, knockout of HAA1 could cause not only statistically significant expression changes of MOT3 knockout affected genes, but also statistically significant expression changes of ECM22 knockout affected genes that are not MOT3 gain-of-regulated genes. Together with the observation above that genes encoding ECM22 and MOT3 were differentially down-expressed when the knockout of HAA1, these results indicated that HAA1 mutant could be used as an approximation to double knockouts of ECM22 and MOT3.

If MOT3 regulated a portion of ECM22 knockout affected genes in the ECM22-deficient cell (i.e. if there exist MOT3 gain-of-regulated genes in the ECM22-deficient cell), knockout of HAA1, which was used as an approximation to double knockouts of ECM22 and MOT3, should reduce MOT3 regulatory effects on MOT3 gain-of-regulated genes according to Figure 1E. Indeed, MOT3 gain-of-regulated genes showed comparable changes in gene expression with the remaining genes upon knockout of HAA1 (P=0.2, Mann-Whitney U-test, Figure 2C, S1).

### Most of apparent-potential TF pairs are cooperative TFs in wild type

Whether apparent TFs and their paired potential TFs have properties in common besides having PPI was investigated. An interesting hypothesis is that these TFs bind target genes in common. Indeed, 13 (65%) of 20 apparent-potential TF pairs are cooperative TFs in wild type: Apparent TF bound genes are statistically significantly bound by the paired-potential TF (P<0.05, chi-square test), and potential TF bound genes are also statistically significantly bound by the paired-apparent TF (P<0.05, chi-square test, Figure 2D, Table S1). It is likely that TF cooperation is just a feature of TFs showing PPI but not a feature of apparent-potential TF pairs. To test this possibility, all TF pairs showing PPI were identified. Only ∼27% of all TF pairs showing PPI are cooperative TFs, which is much lower than the frequency of apparent-potential TF pairs that are cooperative TFs. These results indicate that TF cooperation is a feature of apparent-potential TF pairs. Moreover, among the remaining 7 apparent-potential TF pairs, there are 2 TF pairs in which potential TFs show PPI with TFs that statistically significantly bind genes bound by the paired apparent TFs (P<0.05, chi-square test, ARG81-UME6, MSN4-HSF1). Although these potential TFs do not directly bind genes bound by their paired apparent TFs, it is possible that these potential TFs belong to the protein complexes that regulate these genes, thus these potential TFs may be cooperative TFs of their paired apparent TFs. Apparent TFs and potential TFs may regulate target genes cooperatively through PPI.

### Gain-of-regulated genes show no expression correlation with original regulated genes in wild type

Whether gain-of-regulatory functions of potential TF in their paired apparent TF-deficient cell are associated with its original regulatory functions in wild type was examined. Wild-type expression correlation between potential TF gain-of-regulated genes and genes bound by the corresponding potential TF in wild type are comparable with those between random gene pairs (P=0.18, Mann-Whitney U-test, Figure 3A). Wild-type expression correlation between potential TF gain-of-regulated genes and genes showing differential expression upon knockout of the corresponding potential TF are even lower than those between random gene pairs (P < 10-26, Mann-Whitney U-test, Figure 3B). Moreover, wild-type expression correlation between potential TF gain-of-regulated genes and genes bound by the corresponding paired apparent TF in wild type are also lower than those between random gene pairs (P < 10-7, Mann-Whitney U-test, Figure 3C). These results together demonstrate that either apparent TF or potential TF does not regulate potential TF gain-of-regulated genes in the wild-type cell.

**Figure 3.**
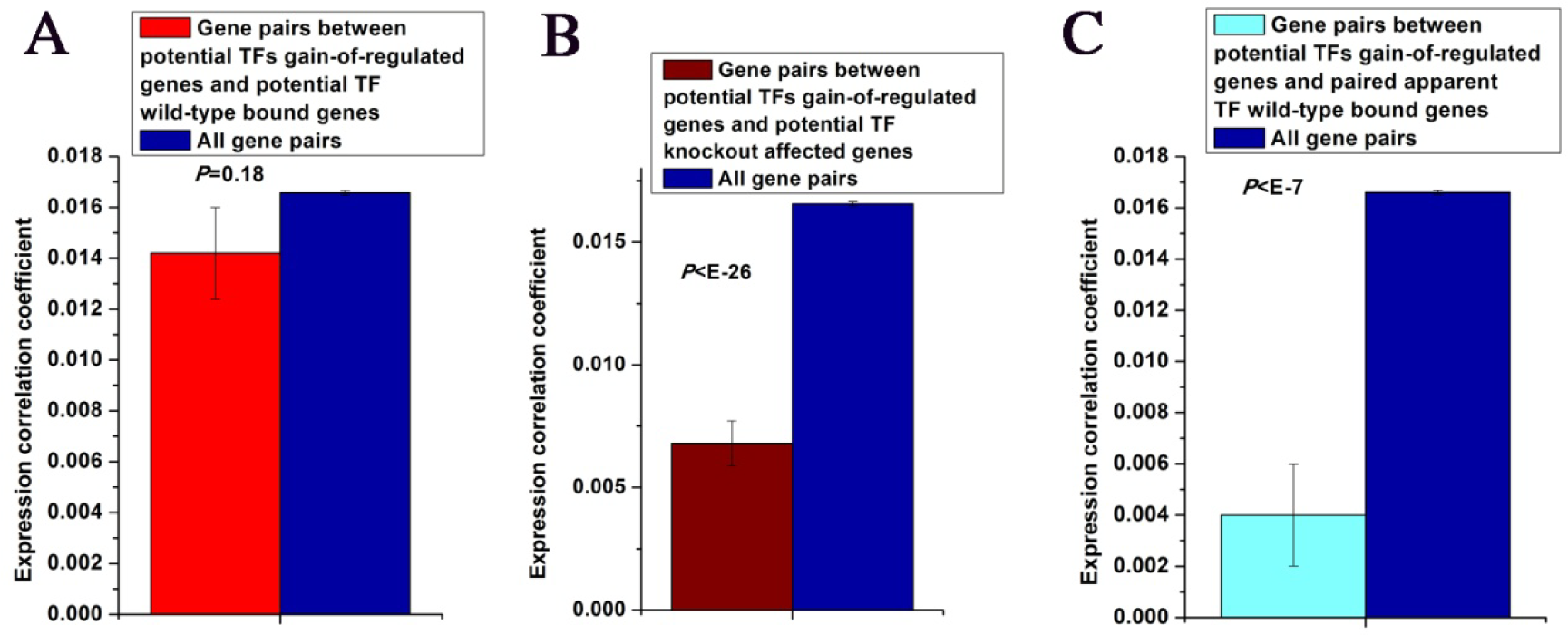
Gain-of-regulated genes show no expression correlation with original regulated genes in wild type. (A) Wild-type Pearson expression correlation between potential TF gain-of-regulated genes and genes bound by the corresponding potential TF in wild type are comparable with those between random gene pairs. (B) Wild-type Pearson expression correlation between potential TF gain-of-regulated genes and genes showing differential expression upon knockout of the corresponding potential TF are lower than those between random gene pairs. (C) Wild-type Pearson expression correlation between potential TF gain-of-regulated genes and genes bound by the corresponding paired apparent TF in wild type are also lower than those between random gene pairs. Error bars were calculated by bootstrapping. The statistical significant values calculated from Mann-Whitney U-test were indicated.

### Potential TF bound genes are affected by knockout of the paired apparent TFs

In my proposed model, potential TFs may gain new regulatory genes in the apparent TFs-deficient cell. Whether this change has effects on potential TFs wild-type regulatory genes was investigated. Whether potential TFs wild-type bound genes show differential expression when knockout of the paired apparent TFs was examined. To avoid confusion, the 13 apparent-potential TF pairs that bind target genes in common were excluded, resulting in 7 apparent-potential TF pairs. 2 of 7 apparent-potential TF pairs show the following property: potential TFs wild-type bound genes showed statistically significant changes in gene expression compared with the remaining genes upon knockout of the paired apparent TFs (P<0.05, Mann-Whitney U-test, Figure 4). This result suggests that knockout of apparent TFs may not only enable their paired potential TFs to bind new targets, but also alter the original binding targets of potential TFs.

**Figure 4.**
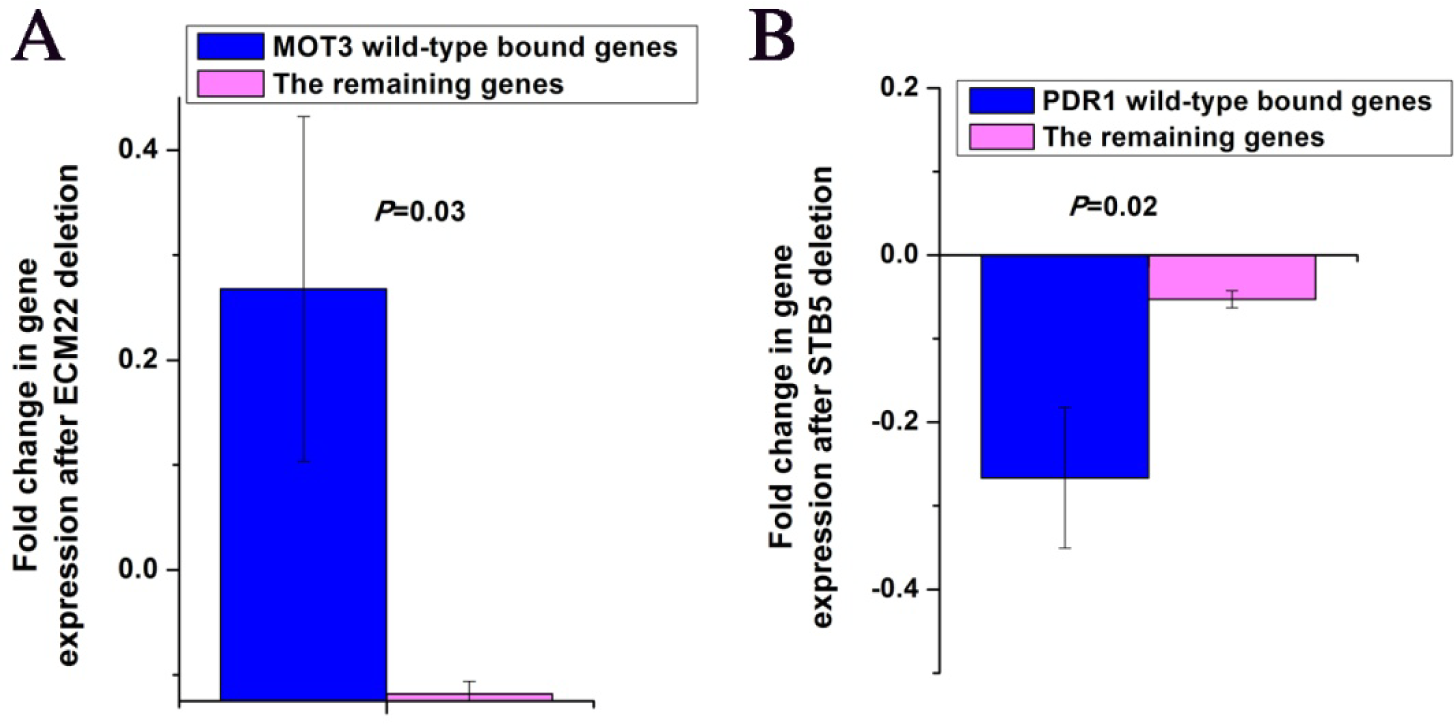
Potential TFs wild-type bound genes showed statistically significant changes in gene expression compared with the remaining genes upon knockout of the paired apparent TFs. (A) ECM22-MOT3 pair. (B) STB5-PDR1 pair. Error bars were calculated by bootstrapping. The statistical significant values calculated from Mann-Whitney U-test were indicated.

## Discussion

TF perturbation can lead to a rewiring of specific transcriptional regulatory circuits, then generate new patterns of gene expression. For example, duplication of a gene coding for a TF can lead to neofunctionalization, where one of the duplicates acquires a novel function that was absent in the pre-duplication protein [22]. In this study, whether the knockout of a TF can have a similar effect was investigated. The knockout of a TF was hypothesized to release another TF, which is the PPI partner of the deleted TF. The free TF can bind new target genes and cause their changes in gene expression. The knockout of a TF may lead to neofunctionalization of another TF.

To examine this hypothesis, multiple sources of data and ten criteria were used to identify 20 such apparent-potential TF pairs in budding yeast. These strict criteria were used to control false positive, meanwhile, some positives might be missed. For example, apparent TF and potential TF should not show PPI with TFs that have binding target enrichment in apparent TF knockout affected genes. If this criterion is abolished, the number of identified TF pairs will be increased to 28. This criterion was used to reduce the possibility that either apparent TF or potential TF is linked to potential TF gain-of-regulatory genes in the wild-type cell, increasing the reliability of my method for identifying apparent-potential TF pairs.

Although I demonstrated that TF-target relationships can be complicated by knockout analysis, I still believed that knockout analysis is a critical method to investigate TF and gene functions. Numerous studies have used this tool to reveal new gene functions successfully. The contribution of this study is to remind researchers to use knockout analysis with caution. It is better to integrate knockout analysis with other datasets generated from other tools to reveal gene functions. For example, it is necessary to integrate TF knockout data, TF-DNA binding data, regulatory cascade data and PPI data to identify TF-target relationships.

Previous studies have tried to explain the small overlap between TF knockout affected genes and TF bound genes from different perspectives, including backup mechanisms in gene regulatory networks [13] and robustness of transcriptional regulatory program [23]. In this study, the small overlap was sought to be explained by proposing a model in which the knockout of a TF can lead to gain-of-regulatory genes of another TF.

I have only begun to explore the transcription factor-target relationships potentially complicated by knockout analysis. To this end, I have proposed a relatively simple model to explain this possibility. There are some limitations in this study. First, there may be other possible models. Another alternative model can be proposed (Figure S2). Multiple sources of data were used to examine this model, but no TFs were found to meet the criteria in the model. It is interesting to propose and validate other models to address the issue. Second, although one my identified pair was validated using other independent data, it will be better to validate more pairs using biological experiments. I would appreciate joining forces with others to further study transcription factor-target relationships potentially complicated by knockout analysis.

