## Supplementary material for "Transcription factor-target relationships complicated by knockout analysis": Figure S1, Figure S2, Table S1

Zhiming Dai[1§]

[1]School of Data and Computer Science, Sun Yat-Sen University, Guangzhou 510006, China

[§]

**Figure S1, S2**
**Table S1**

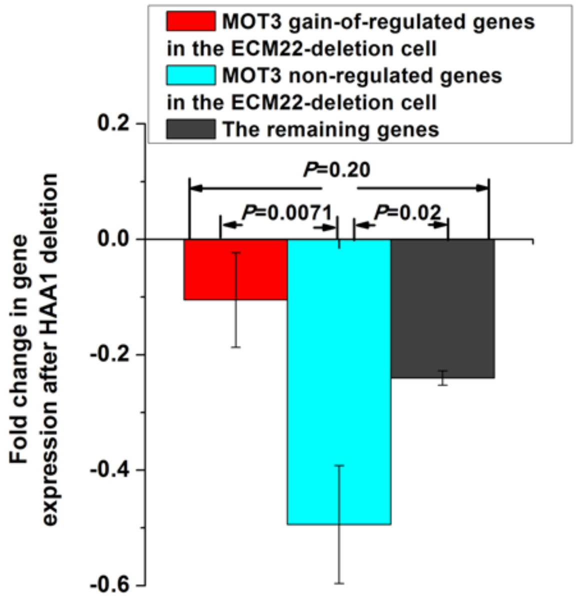


Figure S1. Bars for MOT3 non-regulated genes and MOT3 gain-of-regulated genes in Figure 2C. MOT3 non-regulated genes in the ECM22-deficient cells showed statistically significant changes in gene expression compared with the remaining genes upon knockout of HAA1. MOT3 gain-of-regulated genes in the ECM22-deficient cell showed comparable changes in gene expression with the remaining genes upon knockout of HAA1. Error bars were calculated by bootstrapping. The statistical significant values calculated from Mann-Whitney U-test were indicated.

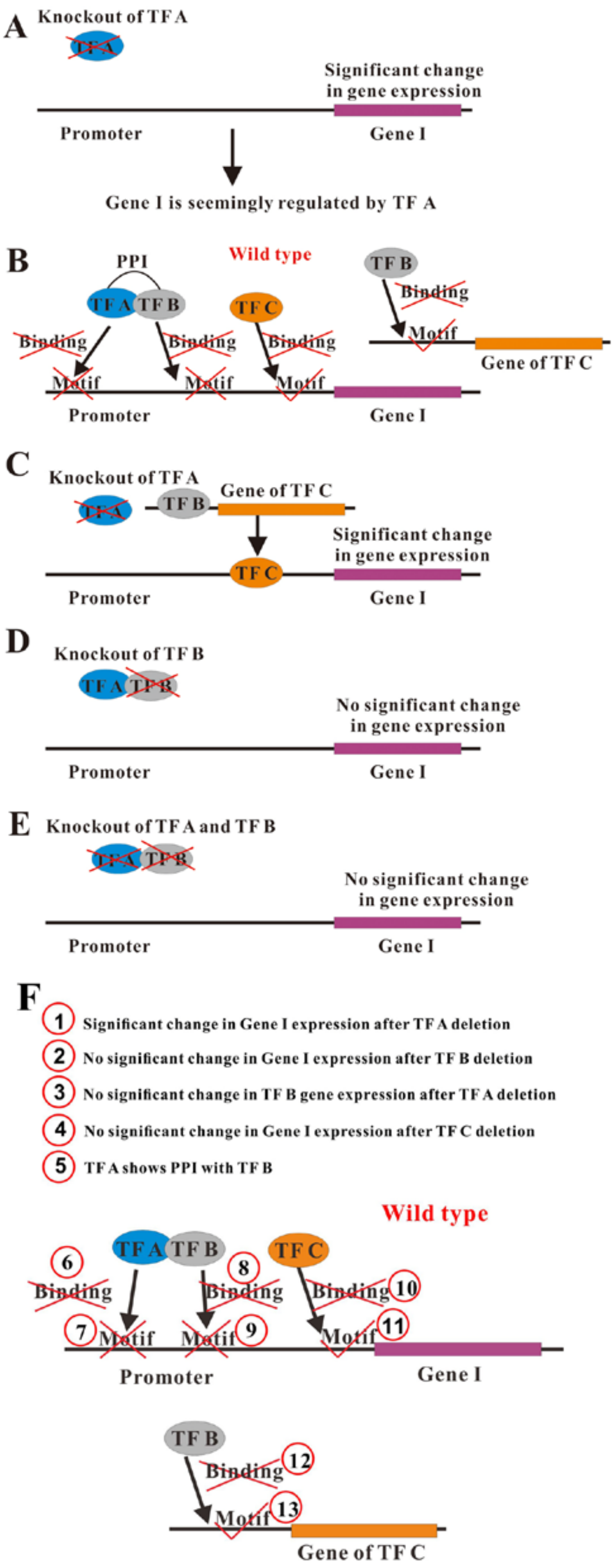


Figure S2. An alternative model for transcription factor-target relationships complicated by knockout analysis. (A), (D) and (E) are the same as Figure 1. (B) Different from Figure 1, Gene I has no TF B bound motifs in promoter region. On the other hand, this gene has DNA motifs for TF C potential binding, though this gene is not bound by TF C in the wild-type cell. The gene coding for TF C has DNA motifs for TF B potential binding, though this gene is not bound by TF B in the wild-type cell. (C) Knockout of TF A may release its PPI partner TF B. The free TF B could find its bound motifs in the gene coding for TF C to up-regulate this gene, though TF B could not bind this gene in wild type due to the limitation imposed by its PPI partner TF A. The up-regulated TF C bind promoter region of Gene I. The so-called TF A regulatory gene (i.e. Gene I) is actually TF C gain-of-regulatory gene in TF A-deficient mutant. This actual regulatory relationship could not be revealed by knockout of TF A. Moreover, This gain-of-regulatory relationship could not be observed by either knockout of the TF B (D) or double knockouts of TF A and TF B (E). (F) The criteria used to identify TF C that meets the description in (B) and (C).

**Table S1 List of apparent-potential TF pairs that have statistically significant overlap in binding target genes.**

| | |
|---|---|
| ARG81 | ARG80 |
| ARG81 | MCM1 |
| HAP5 | HAP3 |
| INO4 | INO2 |
| INO4 | OPI1 |
| INO4 | TYE7 |
| MBP1 | SWI6 |
| MET32 | CBF1 |
| SMP1 | MSN2 |
| SMP1 | MSN4 |
| SMP1 | RLM1 |
| SMP1 | SKO1 |
| STB1 | SWI6 |
